# Digital Light Processing 3D Printing enables High Throughput Fabrication of Human Engineered Heart Tissues for Disease Modeling

**DOI:** 10.1101/2024.10.01.616163

**Authors:** Abhishek P. Dhand, Miranda A. Juarros, Thomas G. Martin, Gabriel J. Rodriguez-Rivera, Dakota R. Hunt, Mackenzie C. Obenreder, Cody O. Crosby, Bianca Meurer-Zeman, Quentin McAfee, Henry Valle-Ayala, Hannah M. Zlotnick, Declan N. Goddard, Christopher C. Ebmeier, Jason A. Burdick, Leslie A. Leinwand

## Abstract

3D *in vitro* engineered heart tissue (EHT) models recapitulate aspects of native cardiac physiology but are often limited by scalability, cost, and reproducibility. Here, we report a simple, one-step method for rapid (∼minutes) fabrication of molds using digital light processing (DLP)-based 3D printing that support the formation of EHTs by human induced pluripotent stem cell derived cardiomyocytes (iPSC-CMs) with high reproducibility (>95% efficiency) and varied designs (e.g., length, aspect ratio). Compared to 2D iPSC-CMs, 3D EHTs display enhanced maturity, including increased expression of β-oxidation genes, higher concentrations of sarcomeric myosins, improved sarcomere density and alignment, and enrichment of cardiac pathways (e.g., upregulation of sodium channels, action potentials, contraction). The technology is applied to model pathological cardiac hypertrophy *in vitro*, using either (i) acute adrenergic agonism or (ii) chronic culture within stiff hydrogel molds. Treated EHTs exhibit increased levels of pathology-associated gene expression and activation of signaling cascades involved in pathological remodeling compared to untreated controls or treated 2D iPSC-CMs. Thus, our method results in robust yet simpler, cheaper, and faster EHTs to study cardiac disease.

## MAIN TEXT

Drug discovery is expensive and time consuming, with many drug candidates failing in the transition from pre-clinical to human clinical trials due to the lack of consideration for species variation and shortcomings of *in vitro* testing^1^. However, growing support from regulatory agencies for the use of *in vitro* models to study human diseases at large scales has increased interest in this area^2^. Additionally, the use of either patient-derived or genome-edited human induced pluripotent stem cells (iPSCs) has enormous benefits for understanding human disease pathogenesis, drug discovery and development, particularly as they overcome the genetic limitations of animal models^3^. Advances in 3-dimensional (3D) *in vitro* platforms also overcome many of the drawbacks to more traditional 2-dimensional (2D) *in vitro* cell cultures, which lack the mechanical environmental cues and 3D complexity of tissues^4^. Specifically, the engineering of 3D cellular systems that confer more of the physiological signatures of tissues in patients will help advance therapeutic discovery and even impact personalized medicine^5^. 3D *in vitro* platforms also enable the manipulation of cellular crosstalk and cell–extracellular matrix (ECM) interactions in a controlled manner, deconvoluting the complexity of signaling cascades present *in vivo*. Importantly, biofabrication methods are advancing to offer a variety of routes to generate complex 3D biological constructs that replicate the functional organization of human tissues, while promoting physiologically relevant cellular interactions^6^.

Heart disease remains the number one cause of death in the world and thus there is an urgent need to better understand the molecular mechanisms involved in cardiac diseases for the development of new therapies^7^. Cell-cell and cell-ECM interactions play an important role in maintaining tissue homeostasis; thus, to study the cardiac response to injury, there is a need to recapitulate this tissue complexity *in vitro*^8^. Current cardiac *in vitro* models include 2D cell monolayers, spheroids, and 3D engineered heart tissues (EHTs)^9,10^. iPSC-derived cardiomyocytes (iPSC-CMs) seeded onto 2D substrates or aggregated in the form of cardiac spheroids can enable high throughput studies but lack the topographical cues and structural alignment of native tissue. Alternatively, cardiac tissues can be engineered through encapsulation of cardiomyocytes in natural or synthetic biomaterials to provide a more appropriate mechanical microenvironment^11–14^. 3D bioprinting can achieve structural and biochemical complexity but is not currently capable of the throughput necessary for industrial or pre-clinical applications, such as drug screening^15–18^. More recently, efforts have focused on designing microphysiological systems such as heart-on-a-chip devices that include multiple cell types (e.g., endothelial cells, fibroblasts, macrophages) with or without electrical/mechanical stimulation^8,19–21^ and have been applied for disease modeling or drug screening^22–26^. These systems can improve the assembly and maturation of tissues *in vitro*, but they require complex setups for execution and are difficult to scale-up, which limits their throughput and widespread adoptability^27^. Thus, there exists a need for a system that exhibits appropriate biological complexity to effectively model human disease, but still can be implemented at a throughput that is useful to the field^28^.

An alternate strategy for the fabrication of EHTs is the seeding of cell-laden ECM gels in devices that provide biophysical cues during tissue assembly, where cell-mediated ECM remodeling causes passive uniaxial tension and induces alignment of myofibrils in cardiomyocytes within microtissues. Existing methods for fabricating such devices include multi-layer soft-lithography or machining^13,29,30^. Some methods rely heavily on multiple rounds of replication with poly(dimethyl siloxane) (PDMS) elastomer or hydrogels to produce features of interest^31–33^. These processes require highly specialized equipment and are time-consuming, costly, limited to simple geometries, and not compatible with large scale manufacturing^34,35^. As an alternative, the use of hydrogels and vat photopolymerization represent an attractive opportunity to overcome these limitations. Specifically, Digital Light Processing (DLP) is an emerging biofabrication technique that enables the 3D printing of materials by crosslinking a liquid resin into a solid hydrogel in a layer-by-layer manner^36,37^. DLP resins exhibit low viscosity and undergo rapid gelation, which results in fast and high throughput fabrication, and DLP-printed hydrogels have been widely adopted for use in tissue repair, delivery of therapeutic cargo, and tissue adhesives, but the use of this modality in 3D *in vitro* models has been limited.

In this article, we leverage DLP-based 3D printing to create hydrogel molds towards the formation of human EHTs. DLP-based 3D printing of hydrogels enables one-step fabrication of micropillar devices with high precision and throughput where the design can be readily altered through simple changes in the computer-aided design (CAD) inputs into the printer. The hydrogel micropillar molds serve as templates to enable the innate self-organizing capability of cells to condense into tissues through the remodeling of surrounding ECM. This simple approach results in morphologically aligned and functionally complex EHTs using a low number of iPSC-CMs and commercially available materials that can be easily implemented across laboratories. Further, our EHT method showed markedly improved metabolic and sarcomeric maturity over 2D *in vitro* cultures with many similarities to mature intact heart tissue and responded to pathological stimuli by displaying expected *in vivo* disease phenotypes that were not found in 2D cultures. Overall, we demonstrate a bottom-up engineering approach for the formation of EHTs that can be employed for robust cardiac disease modeling at large scales with consideration of key design criteria such as throughput, cost and accessibility, as well as robustness and reproducibility.

## RESULTS AND DISCUSSION

### DLP printed hydrogel micropillar molds facilitate formation of uniform EHTs

A resin containing poly(ethylene glycol) diacrylate (PEGDA, macromer) along with the photo-additives LAP (photoinitiator) and tartrazine (photoabsorber) was employed for DLP printing. PEGDA is chemically and biologically inert and thus ensures compatibility with the formation of microtissues. Further, these reagents are commercially available, which allows unrestricted access and reproducibility across batches. The photoinitiator absorbs light to facilitate cleavage to generate free radicals and initiate crosslinking, whereas the photoabsorber provides dose-dependent delays in the initiation of polymerization by competing with photoinitiators to absorb light^38^. The balance of photoinitiator and photoabsorber is integral for successful photolithography-based printing and their absorption maximum should ideally match the wavelength of the light source (**Fig. 1a**). The spectral properties of the light source within the DLP printer were characterized to be 405 ± 20 nm (**Fig. 1b**), which overlaps with the absorption maxima of the chosen photoinitiator (LAP) and photoabsorber (tartrazine) (**Extended Data Fig. 1**). Increasing the photoabsorber concentration reduced the light penetration depth (D_p_) and increased the time (t_g_) required to reach gelation (**Extended Data Fig. 1**).

**Fig 1.**
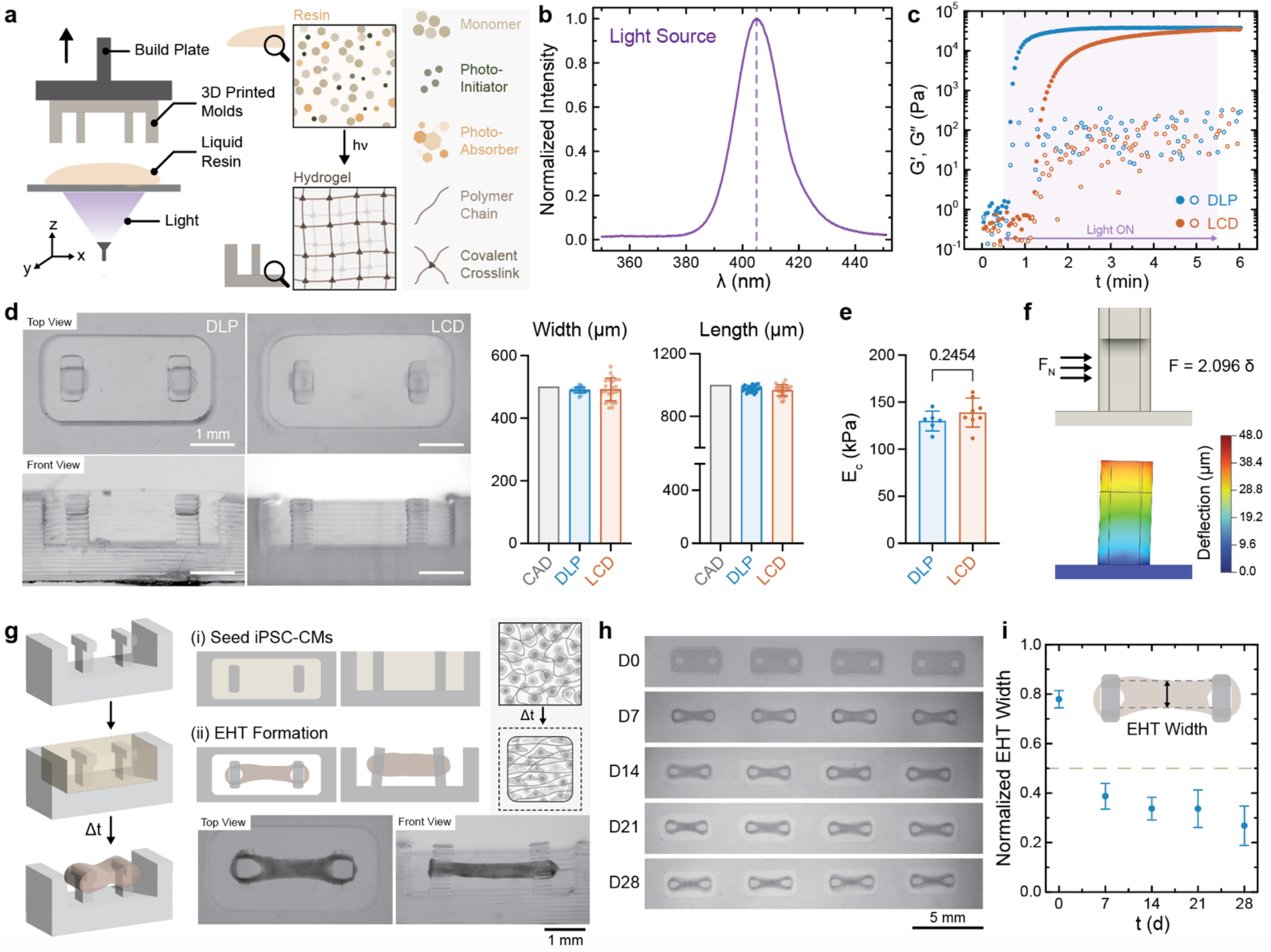
3D printing of hydrogel micropillar molds to fabricate human engineered heart tissue (EHTs). **a**, Schematic representation of digital light processing (DLP) to form 3D micropillar molds using photoreactive resin containing poly(ethylene glycol) diacrylate (macromer), LAP (photoinitiator), and tartrazine (photoabsorber). **b**, Characterization of the spectral properties of the light source within the DLP printer. **c**, Representative photorheology: storage (G′, Pa, closed circles) and loss (G″, Pa, open circles) moduli over time of precursor with varying tartrazine concentration. Shaded area indicates light irradiation. **d**, Photograph (top and cross-sectional front view) of hydrogel micropillar mold 3D printed on a DLP or liquid crystal display (LCD) printer. Scale bars: 1 mm. Comparison of CAD and 3D printed (post-swelling) of micropillar width and length (top view). Data are reported as mean ± SD, n ≥ 20. **e**, Compressive moduli (E_C_, kPa) of 3D printed (with DLP or LCD printer) hydrogel molds after equilibrium swelling. Data are reported as mean ± SD, n ≥ 6, student’s t-test. **f**, Finite element prediction of micropillar deflection in response to static force (F_N_, 100 μN). **g**, Schematic representation of the process flow for EHT formation. iPSC-CMs seeded in an isotropic matrix (day 0) undergo remodeling over time that results in alignment. Representative photographs (top and cross-sectional front view) of fixed EHTs within the micropillar molds at day 28. **h**, Representative images depicting the time course for tissue compaction. Scale bars: 5 mm. **i**, Quantification of width of EHT normalized to the width of the micropillar over time. Data are reported as mean ± SD, n ≥ 18 independent tissues across two biologically independent experiments.

Optimization of the resin formulation (photoinitiator or photoabsorber concentration) is critical to achieve high resolution, especially when printing fine features^39^. Therefore, we evaluated the resolution of hydrogel micropillar molds printed with a traditional DLP printer, as well as with an LCD printer. When compared to DLP printers that use a digital micromirror device (DMD), LCD printers use liquid crystal displays to generate images and are therefore much cheaper. Photorheology was used to investigate the kinetics of gelation of the resin upon light irradiation matching the intensity of the DLP or LCD printer. The time to reach gelation upon light irradiation is marked by the crossover between storage (G′) and loss moduli (G″). The resins described here undergo rapid photo-crosslinking on the order of ∼8 s for the DLP printer, whereas longer exposures (∼44 s) were necessary for the LCD printer owing to the 10-fold lower light intensity (**Fig. 1c**). Further, the dimensions of the printed hydrogels (after equilibrium swelling) matched those defined in the CAD file, with no significant differences in the feature size between those printed on the DLP or LCD printer (**Fig. 1d**). Similarly, the compressive moduli of the DLP printed hydrogels (∼129 kPa) were not statistically different from those fabricated on the LCD printer (∼138 kPa) (**Fig. 1e**). Although the remainder of the work was performed with hydrogels printed on the DLP printer, these findings emphasize the compatibility of the approach with low-cost printers, overcoming cost limitations and maintaining accessibility.

In addition to testing in compression, the tensile modulus of the hydrogels printed on the DLP printer was ∼51 kPa and the storage modulus was ∼45 kPa (**Extended Data Fig. 1**). Despite their high water content (∼86%), 3D printed hydrogels demonstrated high resolution in xy and z-direction (**Extended Data Fig. 1**) and maintained structural stability over time, including in physiological medium at 37°C with no significant changes in the compressive modulus over 35 days (**Supplementary Fig. 1**). The stiffness of the micropillars is important to control the boundary conditions imposed for microtissue formation. To empirically measure the stiffness of the hydrogel micropillars, simulation of pillar deflection in response to varying applied forces was performed. Increased pillar deflection was observed with an increase in applied force, such that the stiffness was estimated to be 2.096 μN μm^-1^ (**Supplementary Fig. 2**). In addition to maximum pillar deflection, local deflection visualized along an individual pillar suggested a gradient in deflection with minimal bending observed closest to the fixed end (**Fig. 1f**).

**Fig 2.**
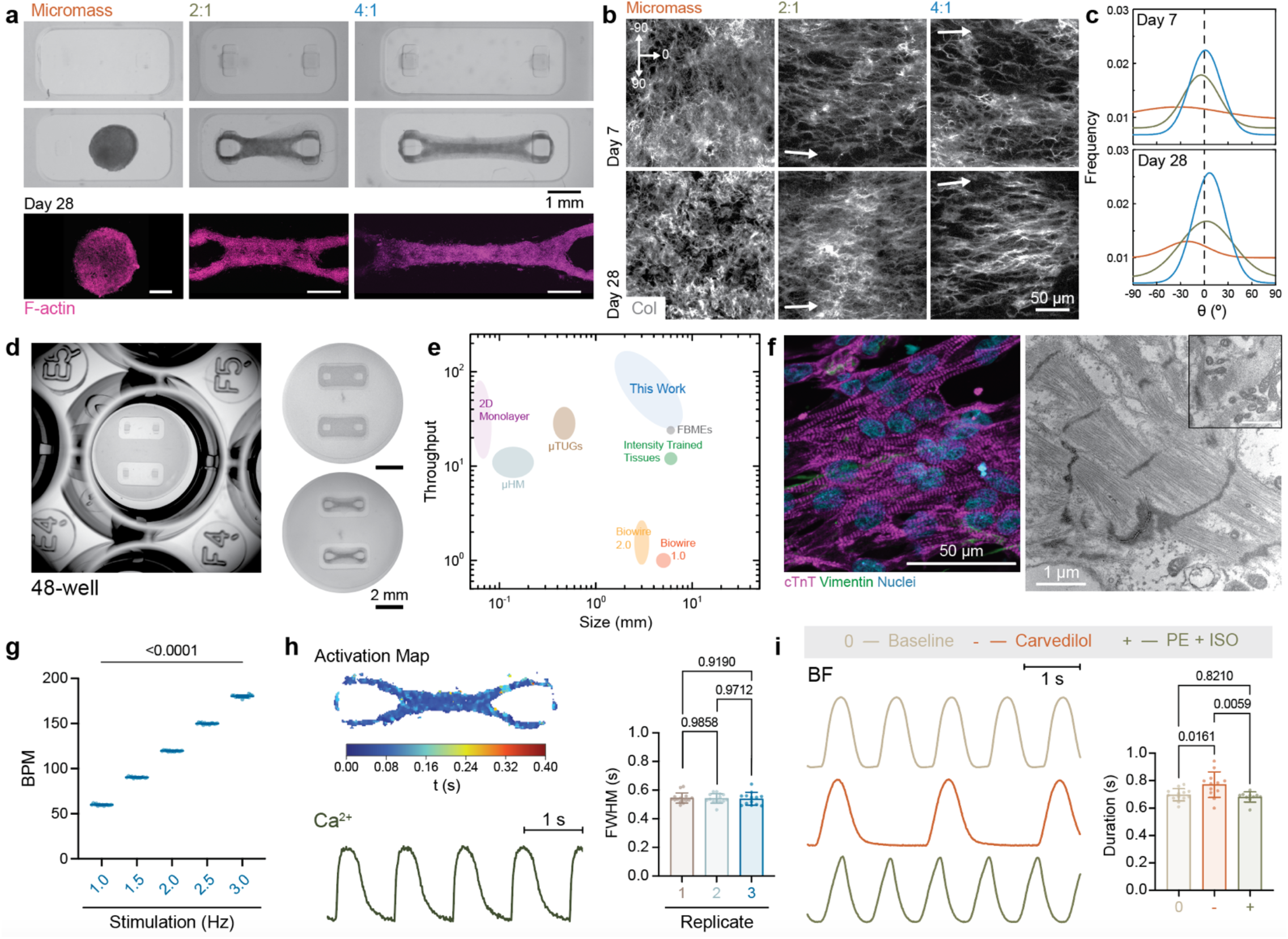
Versatility of DLP enabled formation of EHTs. **a**, Photographs of DLP printed molds with varying distance between micropillars demonstrating ease of design. Representative bright-field and maximum projection fluorescence images of EHTs at day 28. Micromass denotes cell and matrix aggregates formed in molds without micropillars while 2:1 and 4:1 indicates distances of 2 and 4 mm between micropillars, respectively. Scale bars: 1 mm, F-actin (magenta). **b**, Fluorescence images depicting collagen I (gray) remodeling across groups (micromass, 2:1, 4:1) over time. Arrow indicates dominant direction of alignment. Scale bars: 50 μm. **c**, Quantification of collagen I alignment (from b) as a function of angle (8, °). **d**, Photograph of DLP printed hydrogel micropillar mold adapted to fit in a 48-well format. Representative images of EHTs at day 0 and day 28. Scale bars: 2 mm. **e**, Comparison of EHT size and throughput of production of previously reported studies with those reported in the present work. **f**, Fluorescence image of EHT immunostained for cardiac troponin T (cTnT, magenta), vimentin (green), and cell nuclei (Hoechst 33342, cyan) at day 28. Transmission electron microscopy (TEM) of EHTs at day 28 depicting sarcomeres. Scale bars: 2 μm. **g**, Beating rate (beats per minute, BPM) of EHTs in response to electrical stimulation with increasing frequency at day 28. Data are reported as mean ± SD, n ≥ 12 independent tissues, one-way ANOVA followed by Tukey’s post-hoc test. **h**, Activation map and calcium trace of EHTs during field stimulation (1 Hz) at day 28. Quantification of full width half maximum (FWHM) for calcium traces. Data are reported as mean ± SD, n ≥ 12 independent tissues across three biologically independent experiments, one-way ANOVA followed by Tukey’s post-hoc test. **i**, Contraction profile and contraction duration of EHTs at baseline and after treatment with carvedilol and PE+ISO (phenylephrine, isoproterenol) at day 28. BF denotes bright field imaging. Data are reported as mean ± SD, n ≥ 10 independent tissues, one-way ANOVA followed by Tukey’s post-hoc test.

Upon optical and mechanical characterization, hydrogel micropillar molds were next employed to form EHTs. Collagen I is the primary structural ECM protein in the heart that supports alignment and function, transfers the force generated by cardiomyocytes, and provides passive tension during diastole^40^. Hence, we selected collagen I as the main matrix component for the EHTs. Collagen supplemented with ECM factors (i.e., Matrigel® basement membrane) and containing iPSC-CMs was pipetted into the micropillar molds (denoted as day 0). Over time, the cells remodeled the surrounding matrix and compacted the collagen gels as evidenced by the dramatic reduction in tissue width normalized to the micropillar width by day 7 and reached a plateau by day 14 after seeding (**Fig. 1g-i**). The dynamic interplay between cells and ECM can be observed as the cells synthesize and remodel their surrounding matrix. Moreover, the formation of EHTs between hydrogel micropillars allows the developing tissues to perform contractile work against the resistance imparted by elastic pillars, similar to the *in vivo* physiology of strain imparted by volume changes in the ventricle. EHTs displayed synchronous and spontaneous beating within the first week of culture as evidenced by the deflection of micropillars. Variability in the efficiency of tissue formation and the range of tissue sizes can be considerable limiting factors across methods^27,29^. Therefore, it is important to note that the size of the microtissues in this study was homogeneous across multiple, independent replicates, while the efficiency of formation of these microtissues was ∼96% with our DLP printed micropillar molds as compared to roughly 50% efficiency with previously reported miniaturized tissues^29^ (**Supplementary Fig. 3**).

**Fig 3.**
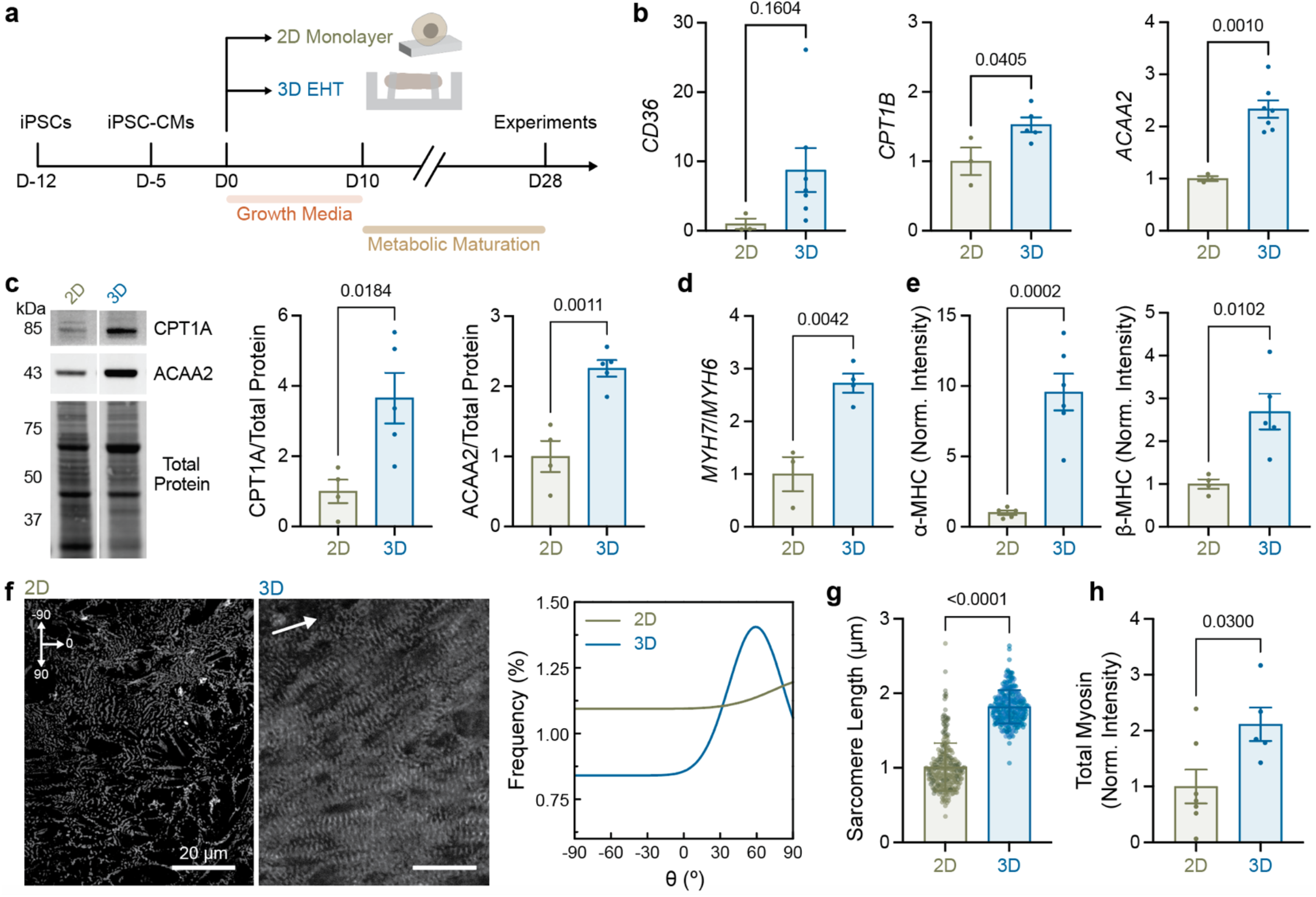
3D EHTs display improved metabolic and sarcomeric maturity compared to 2D iPSC-CMs. **a**, Experimental design: human iPSC-CMs seeded in 2D or encapsulated in collagen hydrogel to form 3D EHTs within DLP printed hydrogel micropillar molds and cultured over time. **b**, Relative gene expression of *CD36, CPT1B, ACAA2* normalized to *GAPDH*. Data are reported as mean ± SEM, n≥3 biologically independent experiments, unpaired student’s t-test. **c**, Western blot analysis and relative expression of p-CPT1A and p-ACAA2 normalized to total protein. Data are reported as mean ± SEM, n≥4 biologically independent experiments, unpaired student’s t-test. **d**, Relative gene expression of *MYH7/MYH6*. Data are reported as mean ± SEM, n≥3 biologically independent experiments, unpaired student’s t-test. **e**, Quantification of relative intensity of α-MHC and β-MHC. Data are reported as mean ± SEM, n≥4 biologically independent experiments, unpaired student’s t-test. **f**, Representative fluorescence images and quantification of sarcomere alignment (frequency, %) for 2D iPSC-CMs and 3D EHTs immunostained for α-actinin (gray). Arrow indicates dominant direction of alignment. Scale bars: 20 μm. **g**, Quantification of sarcomere length and **h**, relative myosin intensity. Data are reported as mean ± SEM, n≥4 biologically independent experiments, unpaired student’s t-test.

We next evaluated the influence of cell seeding density (i.e., 10 million cells per mL compared to 60 million cells per mL) and ECM component (i.e., collagen I compared to fibrin) on EHT function (**Supplementary Fig. 4**). Although EHTs with higher cell seeding density exhibited longer relaxation times, no significant differences were observed in force generation. Hence, to avoid the use of higher number of cells per EHT, cell density of 10 million cells per mL was chosen. Similarly, EHTs with collagen as the primary component generated significantly higher contractile forces when compared to fibrin. This optimization based on functional analysis established the suitability for EHTs with collagen and low cell seeding density for all further experiments in this study.

**Fig 4.**
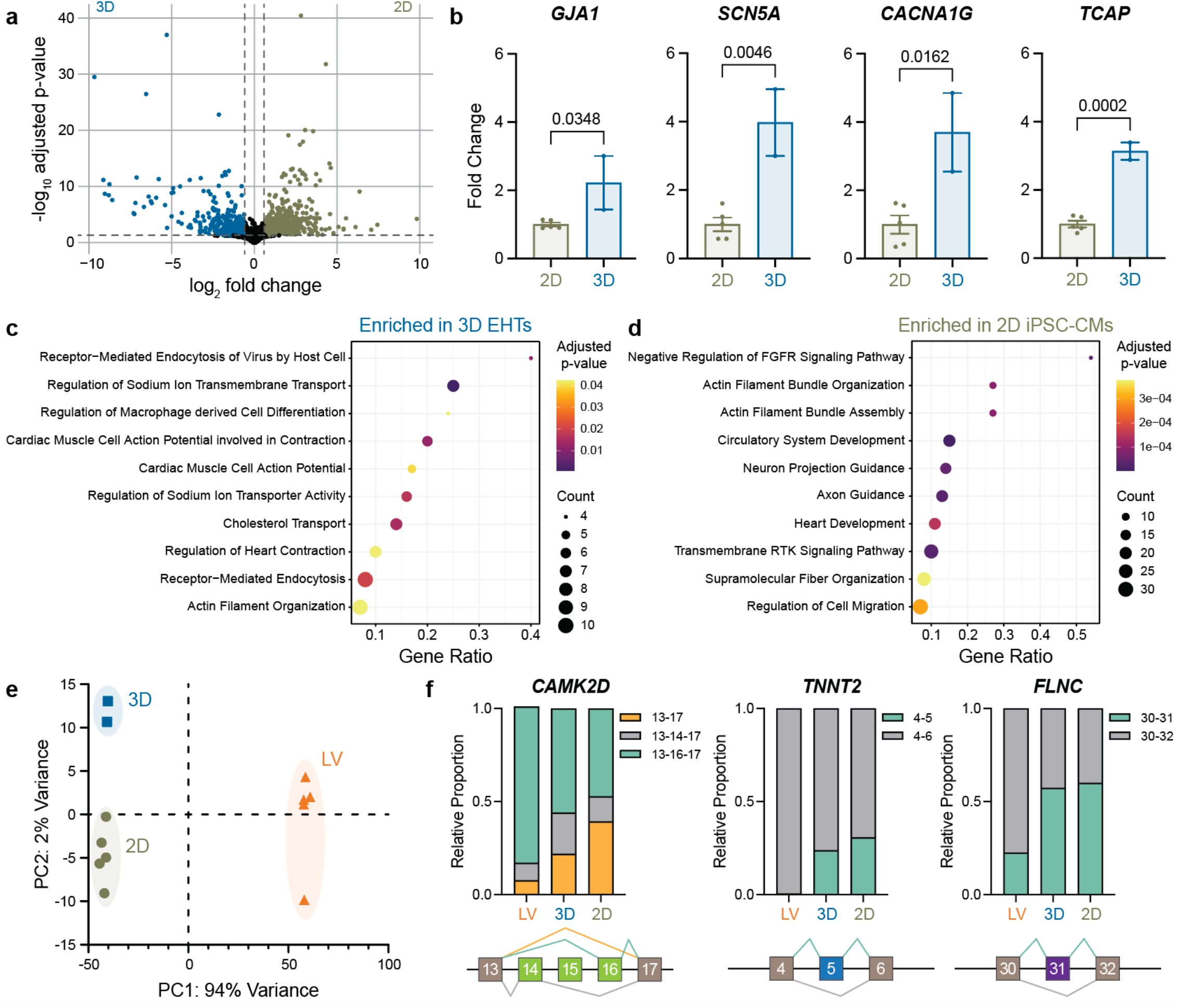
3D EHTs are enriched with ion handling and cardiac contractility genes. **a**, Analysis of genes differentially expressed between 2D iPSC-CMs and 3D EHTs. **b**, Fold change in normalized counts of representative genes *GJA1, SCN5A, CACNA1G*, and *TCAP*. Data are reported as mean ± SEM, n≥2 biologically independent experiments, unpaired student’s t-test. **c**, Genes exclusively upregulated in 3D EHTs analyzed for pathway enrichment. **d**, Genes exclusively upregulated in 2D iPSC-CMs analyzed for pathway enrichment. Results are shown as a bubble plot where bubble color and size represent adjusted p-value and gene count, respectively. **e**, Principal component analyses indicating distinct clustering of 2D iPSC-CMs, 3D EHTs, and non-failing human left ventricle (LV) samples (n≥2 per group). **f**, Analysis of relative proportion of alternate exon splicing isoforms for various genes *CAMK2D* (exons 13-17), *TNNT2* (exon 5), *FLNC* (exon 31) across 2D, 3D, and LV samples.

### DLP printing provides rapid design flexibility towards formation of EHTs

Beyond the ease of micropillar mold fabrication with DLP, the geometric freedom afforded by 3D printing allows for ease in changing the design of the printed object. For instance, iterations of mold design can be conceived, modeled, and 3D printed within a few hours, avoiding logistical constraints (e.g., time, material) of multi-step casting. We demonstrated this by creating hydrogel molds in different shapes (e.g., rectangular pillars, circular pillars, rectangular slabs) and with various modes of tissue anchoring, inspired by previous work (**Extended Data Fig. 2**)^12,41–43^. To further illustrate the design flexibility, molds either without micropillars or with varying distance between micropillars (e.g., aspect ratios of 2:1, 4:1) were fabricated to control the microtissue aspect ratio, as visualized through bright field and F-actin staining of cells within the resulting EHTs (**Fig. 2a**). In the absence of micropillars, the collagen gels underwent isotropic compaction into cell-matrix aggregates (i.e., micromass) due to the absence of passive tension (**Fig. 2a**). As a result, the collagen within these micromass tissues was randomly organized **(Fig. 2b**). In contrast, in the case of both 2:1 and 4:1 aspect ratio molds, the micropillars provided passive tension, which induced alignment of the collagen as early as day 7 as it underwent compaction due to cell-mediated remodeling (**Fig. 2b**). A higher degree of collagen orientation along the longitudinal axis was observed with 4:1 molds when compared to 2:1 molds (**Fig. 2c**), a finding that is consistent with previous work^44^. Thus, DLP printing provides a simple method for fabricating molds that enable collagen alignment within EHTs. To further illustrate design flexibility, we also fabricated micropillar molds that were scaled down to fit within a 48-well plate format, which is highly desirable for screening and discovery applications (**Fig. 2d, Supplementary Fig. 5**). Beyond the context of experimental throughput, formation of EHTs in a 24-or 48-well format provides several other advantages including (i) higher degree of standardization; (ii) repeated, real-time monitoring without compromising mold sterility; (iii) low number of cells required per EHT; and (iv) ability to integrate long-term electrical stimulation or custom-made hardware to automatically measure functional outputs (e.g., beat frequency, contractile force)^45^.

**Fig 5.**
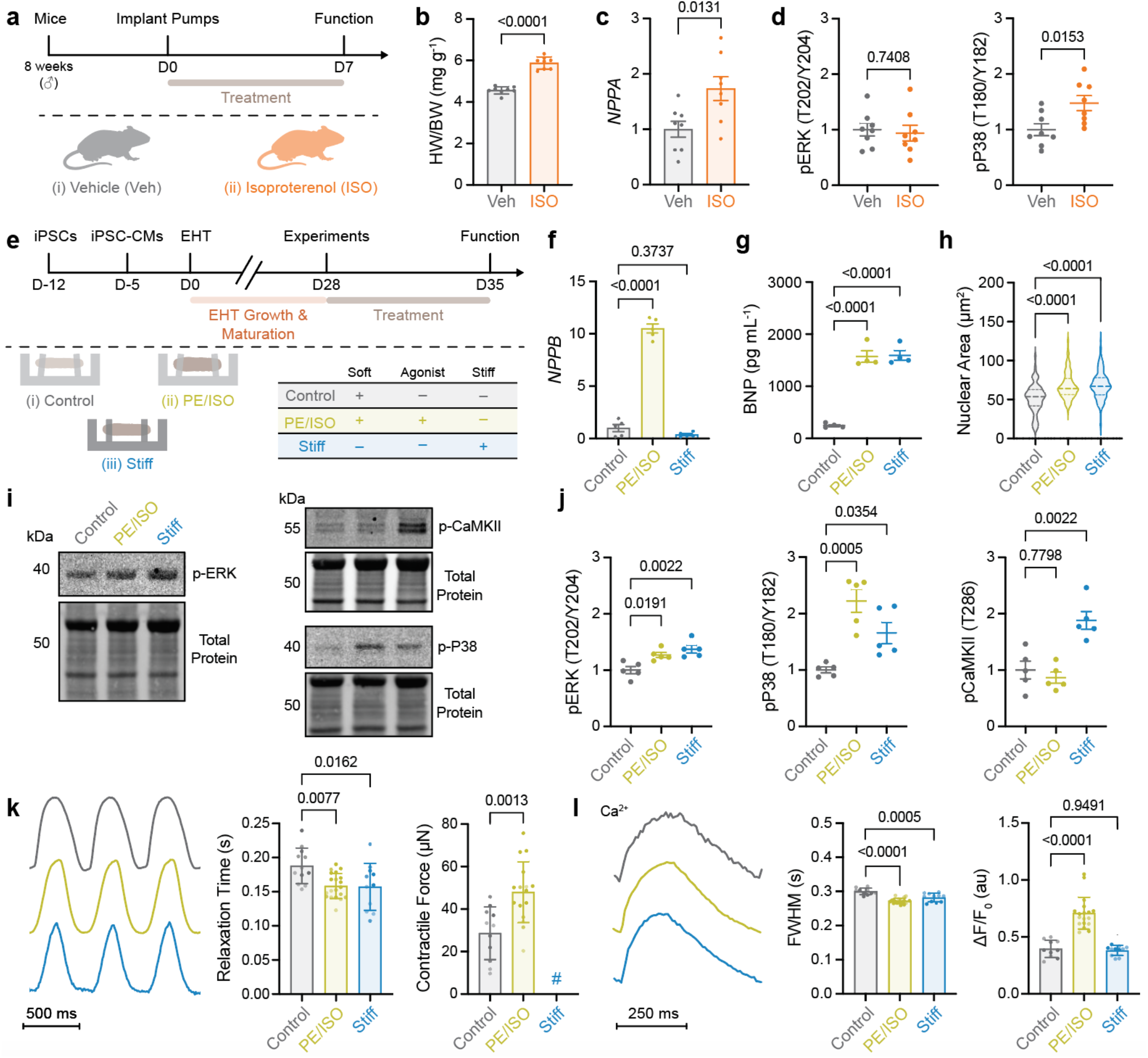
3D EHTs effectively model molecular and functional features of pathology associated with adrenergic agonism and increased afterload. **a**, Experimental design: in vivo implantation of pumps to deliver vehicle or isoproterenol (ISO) in mice. **b**, Comparison of heart weight to body weight of mice after treatment (vehicle vs ISO). **c**, Relative gene expression of *NPPA* normalized to *GAPDH*. **d**, Western blot analysis and relative expression of p-ERK and p-P38 normalized to total protein. Data are reported as mean ± SEM, n≥8 biologically independent experiments, unpaired student’s t-test. **e**, Experimental design: EHTs cultured in soft hydrogel micropillar molds without (C: control) or with adrenergic agonist (phenylephrine, PE and isoproterenol, ISO) treatment (A: agonist) or cultured in stiff hydrogel micropillar molds without agonist (L: stiffness overload). **f**, Relative gene expression of *NPPB* normalized to *GAPDH*; **g**, Measurement of secreted BNP (pg mL^-1^); **h**, Quantification of nuclear area (μm^2^); and **i**, Western blot analysis and **j**, relative expression of p-ERK, p-P38, and p-CamKII normalized to total protein for control, agonist, and stiffness overload treated EHTs at day 29. Data are reported as mean ± SEM, n≥20 tissues pooled each time, N≥4 biologically independent experiments, one-way ANOVA followed by Tukey’s post-hoc test, comparisons of all groups to only control shown on graphs. **k**, Representative contraction profile, relaxation time (s), and contractile force (μN) for control and treated EHTs. # indicates inability to measure force due to minimal deflection of stiff micropillars. **l**, Representative calcium trace, full width half maximum (FWHM, s), and contraction amplitude (ΔF/F_0_, au) for control and treated EHTs. Functional analysis was performed under field stimulation at 2 Hz at day 35. Data are reported as mean ± SD, n ≥ 10 tissues from three independent experiments, one-way ANOVA followed by Tukey’s post-hoc test, comparisons of all groups to only control shown on graphs.

To demonstrate the high-throughput capacity of our methodology, we printed 28 micropillar molds in less than 25 minutes on a build plate measuring 16.5 × 7.2 cm (**Supplementary Fig. 5**). Moreover, the speed of fabrication with DLP is dependent on the height of the object rather than the size in the x-y direction. Therefore, a larger build plate would decrease the time necessary to print a single mold and thus increase the throughput. To place the current study in the context of previously reported EHTs^11–14,29,30,42^, we compared the throughput of fabrication to the size of the EHTs formed (**Fig. 2e**). Smaller EHTs reduce the number of iPSC-CMs required per EHT (e.g., as low as 500 cells) as compared to larger tissues that can require up to 1 to 2 million cells. While miniaturization of EHTs significantly reduces the number of cells and associated cost with iPSC differentiation, manual handling (i.e., casting, media changes) of micro-EHTs is particularly challenging^11,29^. Further, the low amount of RNA or protein recovered from an individual EHT reduces the possibilities for molecular biology or biochemical analyses. In contrast, macroscopic EHTs are easier to handle and can generate forces that can be directly measured with force transducers. Our approach combines the benefits of both microscopic and macroscopic EHTs, wherein the tissues are large enough to produce robust contractions while possessing an anisotropic form. Further, this method requires only 50,000-70,000 cells per EHT, forming tissues with lengths ranging from 2 to 6 mm afforded by the design flexibility discussed earlier. The relatively small size of the EHTs permits real time monitoring and imaging via confocal microscopy to acquire data to assess contractile function.

EHTs displayed a high cardiomyocyte density when immunostained for the cardiomyocyte-specific protein troponin T (cTnT) and the non-myocyte marker, vimentin. Transmission electron microscopy (TEM) of EHTs revealed an orderly register of sarcomeres containing tightly demarcated z-disks within myofibrils, and a high density of mitochondria (**Fig. 2f, Supplementary Fig. 6**). Within immature cardiomyocytes, the mitochondria are randomly distributed and lack well-formed ultrastructure^46^. In contrast, mitochondria within our 3D EHTs occupy a sizable volume fraction of the cell (∼13%) and exhibit well-formed cristae. In addition to examination of cardiac ultrastructure, we measured auxotonic contractile function and calcium handling of EHTs via optical tracking. EHTs were highly responsive to field stimulation up to at least 3 Hz, suggesting strong electrical coupling between cells, likely as a result of improved gap junction formation (**Fig. 2g**)^47^. Increased beat rate upon increasing electrical stimulation frequency was coupled with a subsequent decrease in contraction and relaxation times (**Supplementary Fig. 7**). Contractile force generation is dependent on the interplay between electrical activation, calcium handling, and myofilament activation^48^. Additionally, in the case of EHTs, the force generated can vary as a function of cell purity, cell age, and culture conditions and depends highly on the tissue geometry, local alignment, and tissue stiffness. The EHTs generate about 0.13 to 0.25 mN mm^-2^ contractile stress under 1 Hz stimulation. Although these values are much lower than that reported for adult human ventricular cardiomyocytes, they match well with the other micropillar based systems^49,50^.

Given that calcium plays an integral role in cardiac excitation-contraction coupling, we next assessed the calcium handling capability of EHTs. Upon field stimulation, EHTs displayed high spatial homogeneity in activation delay as visualized through the activation map (**Fig. 2h**). To investigate the reproducibility of formation and function of EHTs, the full width half maximum (FWHM) was analysed from calcium intensity measurements (**Fig. 2h**). No significant differences in FWHM were observed across biologically independent batches, indicating a high level of reproducibility of healthy tissue function enabled by DLP printed hydrogel micropillar molds. While reproducibility plays an important role for these EHTs to be used in a pharmacological setting, another critical feature is their applicability for drug discovery. Our EHTs responded acutely to the α/β-adrenergic receptor blocker carvedilol, as demonstrated through increased contraction duration; however, when treated with positive inotropic drugs (PE/ISO) the contraction duration returned to baseline levels (**Fig. 2i**). In combination with the scalability and limited batch-to-batch variability already described, this finding suggests that these EHTs will be a valuable platform for medium to high-throughput drug discovery.

The versatility and rapid customizability provided by our platform allows for incorporation of a wide variety of cells for fabrication of more complicated geometries readily. Taken together, DLP printing of hydrogel micropillar molds presents a powerful bioengineering toolbox for modeling of cardiac diseases and screening of therapeutics for regeneration.

### EHTs display improved metabolic and sarcomeric maturity over 2D-cultured iPSC-CMs

Maturation of cardiomyocytes *in vivo* in terms of size, shape, metabolism, and physiological function takes years^46^. Therefore, several recent strategies have focused on maturation of iPSC-CMs with long term culture, metabolic cues, and/or mechanical or electrical stimulation^48^. During development, the metabolism switches from glycolysis (fetal) to oxidative phosphorylation (adult)^51^. To consider this, we first compared the effect of culturing EHTs in maturation medium (MM) containing low glucose and supplemented with fatty acids (oleic acid, palmitic acid) and galactose as opposed to traditional growth medium (GM) (**Extended Data Fig. 3**)^52–54^. While there were no significant differences in cell viability between the two conditions, EHTs cultured in MM possessed higher widths and thicknesses compared to those in GM. Further, increased fatty acid uptake and high density of mitochondria were observed for EHTs cultured in MM (**Extended Data Fig. 3**), which likely contributes to improved maturity. Spontaneous beat frequency of EHTs in MM was consistently higher than GM over the period of 35 days (**Extended Data Fig. 3**). Despite the differences in biochemical output, no significant differences were recorded in function (relaxation time, FWHM, or contractile force) (**Extended Data Fig. 3**). Based on these results, subsequent studies were performed in MM.

We next evaluated the degree of maturity of the EHTs compared to standard 2D cultured iPSC-CMs. After iPSCs were differentiated into cardiomyocytes and replated onto 2D tissue culture dishes or the EHT molds, they were switched to MM to promote metabolic maturity (**Fig. 3a**). To assess metabolic maturity of CMs in the two settings, we first measured gene and protein expression of factors involved in fatty-acid transport and oxidation. Compared to 2D iPSC-CMs, we found that EHTs had significantly increased expression of the fatty acid transporters CD36 and CPT1A/B and the fatty acid beta oxidation enzyme ACAA2 (**Fig. 3b-c**). Relative levels of the two major myosin heavy chain (MyHC) isoforms can also be used as an indicator for the level of CM maturity^55^. MyHC is the primary component of the sarcomere thick filament and is the molecular driver of muscle contraction through ATP hydrolysis-dependent displacement of actin filaments by myosin motor domains^56^. Compared with embryonic cells, adult human cardiomyocytes have increased β-MHC (a catalytically slower motor encoded by the *MYH7* gene) and reduced α-MHC (*MYH6*)^55^. We found that EHTs had an increased *MYH7* to *MYH6* gene expression ratio compared to 2D iPSC-CMs (**Fig. 3d**), indicating there was a shift in the EHTs toward expressing a more mature myosin profile at the transcript level. EHTs also displayed increased relative protein abundance of both α-MHC and β-MHC as determined by mass spectrometry analysis compared to 2D iPSC-CMs (**Fig. 3e**), which suggests that there is increased sarcomere expression in the EHTs. During maturation, the sarcomere becomes highly organized, and sarcomere length increases to support force generation. To further assess contractile maturity, we compared sarcomere alignment and length between the two conditions. By orientation analysis, we found that EHTs displayed improved sarcomere organization, alignment, and density compared to 2D iPSC-CMs (**Fig. 3f-h**). In addition, sarcomere length in EHTs averaged ∼1.82 µm, which is similar to the length in relaxed adult cardiomyocytes (1.8 to 2.0 µm) (**Fig. 3g**)^57^. These collective data indicate that the EHT model represents a significant improvement compared to 2D cultures with respect to cardiomyocyte maturity, at the levels of both metabolism and molecular contractile machinery.

Next, to explore global transcriptome differences between EHTs and 2D iPSC-CMs using an unbiased approach, we performed bulk mRNA sequencing (**Fig. 4**). Our analysis identified that genes associated with calcium, sodium, and potassium ion handling (e.g., *ATP1B1, CACNA1G, FXYD1, GJA1, SCN5A*), fatty acid transport (*CD36*), and the sarcomere (*LMOD3, TCAP*, and *TNNI3K)* were among the most differentially expressed between EHTs and 2D iPSC-CMs (**Fig. 4a-b, Supplementary Fig. 8**). The enriched expression of ion handling genes in 3D supports the conclusion that these microtissues have improved electrophysiological properties over 2D cultures, which likely underlies their responsiveness to field stimulation observed above (**Fig. 2g**). To further determine differences between these two conditions, we conducted Gene Ontology Biological Process enrichment analysis. We found that EHTs had increased expression of genes associated with ion transport and heart contraction, while 2D cultures had higher abundance of genes associated with heart development and the actin cytoskeleton (**Fig. 4c-d**). These RNA-seq findings support that there is broad remodeling of the transcriptome in microtissues, which likely contributes to their improved maturity. It is also important to place these data in the context of human cardiac tissue; therefore, we compared the RNA-seq data of EHTs with that from bulk mRNA sequencing on five non-failing human heart left ventricular (LV) samples. While we found that our 2D and 3D EHT conditions were clearly distinct from each other, the EHTs still displayed notable differences in gene expression patterns from the adult human LV (**Fig. 4e**). These differences are expected to be at least partially due to the limited cell type complexity of our model, which is predominantly cardiomyocytes, compared with the human LV, which contains a diversity of cell types^58^. To further examine cardiomyocyte maturity in our EHT model, we also analysed alternative exon inclusion of several cardiomyocyte gene transcripts that change with cardiac maturation. Alternative pre-mRNA splicing can generate multiple unique proteoforms from the same gene product, thereby increasing proteome complexity by an order of magnitude^59^. Switching from embryonic to adult isoforms is known to contribute to cardiomyocyte functional maturation^60^. We assessed alternative exon expression for *CAMK2D* (encoding Ca^2+^/calmodulin-dependent protein kinase IIδ) exons 13-17, *TNNT2* (cardiac troponin T) exon 5, and *FLNC* (filamin C) exon 31 in the RNA-seq datasets, as these splicing events have been well characterized previously^61–63^. In each case, while the EHTs showed a slight trend toward the adult-like splicing pattern, they were still much more similar to 2D than to the human LV (**Fig. 4f**). Thus, our findings indicate that – despite the improved metabolic and sarcomeric maturity detailed above – the EHTs still maintain the immature transcript splicing patterns that are inherent to iPSC-CMs. This is the first study to our knowledge to compare alternative splicing head-to-head in iPSC-CMs and EHTs. Future studies assessing the potential to initiate adult-like alternative splicing in iPSC-CM models will be extremely valuable to further improve their functional maturity.

### 3D EHTs effectively model molecular features of human heart disease

The heart experiences dynamic mechanical forces *in vivo*. These include stretch on the left ventricular tissue during diastole as the chamber fills with blood (preload) and afterload forces imparted by systemic blood pressure on the aortic valve, which the muscle must work against to eject blood during systole. *In vitro*, the preload force can be modeled as the load applied to the EHT prior to contraction, whereas afterload is the resistance experienced by the EHT during contraction^64^. During development, both preload and afterload play a fundamental role in maturation of the contractile machinery and regulation of contractile function. However, in disease states such as pathological cardiac hypertrophy, chronic afterload can result from increased stiffness or increased hemodynamic load due to dysregulation of systemic neurohormonal signaling and/or stenotic remodeling of the aortic valve, all of which may lead to heart failure^7,65^.

Modeling of pathological cardiac hypertrophy in preclinical animal models is commonly achieved through transverse aortic constriction (TAC) or through implantation of mini-osmotic pumps to deliver neurohormonal agonists^7^. These methods can result in induction of robust pathological hypertrophy that is associated with reactivation of the fetal gene program. We employed a common pre-clinical approach to studying heart disease by exposing mice to the pathological stressor, and β-adrenergic agonist, isoproterenol (ISO), which was delivered by a subscapular implanted osmotic minipump (**Fig. 5a**). As expected, these mice developed robust cardiac hypertrophy after one week, as indicated by increased heart weight to body weight ratio (**Fig. 5b**). Pathological cardiac remodeling in this model coincided with induction of molecular features associated with heart disease, including increased atrial natriuretic peptide (*NPPA*) gene expression and activation of P38 MAPK signaling^65^ (**Fig. 5c-d**). Rodent preclinical models are valuable tools for disease modeling; however, such models are limited by their low-throughput, high variability, high costs, ethical concerns, and lack of human physiology. In addition, bulk analyses of heart tissues at the molecular scale can be difficult to interpret due to the presence of non-myocytes^26^. Hence, there is a need for simplified, high-throughput models to mechanistically probe the direct effect of pathological stressors on cardiomyocytes.

To overcome these limitations, we employed our technology to model chronic afterload using two different approaches: 1. A combined treatment with the adrenergic agonists ISO and phenylephrine (PE) (**Fig. 5e**); the choice of using both an β-adrenergic agonist and α-adrenergic agonist in combination was undertaken because these cells express both receptors and EHTs were noted to respond to both PE and ISO alone (**Supplementary Fig. 9**), whereas fully adult cardiomyocytes are responsive only to β-adrenergic agonism; 2. Increasing the EHT mold stiffness (∼27 μN μm^-1^) to model the increased stiffness that occurs in the diseased heart (**Fig. 5e**). Previous studies have reported enhanced afterload via increases in stiffness, insertion of metal braces or magnetics-based approaches^64,66,67^. However, these methods require specialized equipment that may not be commonly available and metal braces exert supraphysiological stiffnesses, which limits their utility in modeling disease. In contrast, our approach relies on the versatility of DLP to 3D print hydrogel micropillar molds with varying stiffness. Tuning the concentration of monomer within the resin provides a simple method for altering the mechanical properties of the micropillars without the need to change the design or use of additional accessories.

To assess the extent of pathology in these conditions, we examined natriuretic peptide expression, as these fetal gene products are canonical markers of pathology and positively correlate with the increased hemodynamic strain observed in heart disease^7^. In addition to metabolic cues, the duration of culture also plays a critical role in EHT maturity. EHTs cultured in MM for shorter duration (i.e., 7 days) were unable to respond to agonist treatment as seen from marginal changes in expression of *NPPB* compared to untreated controls (**Supplementary Fig. 10**). Further, these tissues failed to capture functional (e.g., relaxation time, FWHM) and molecular (e.g., p-CamKII) hallmarks of pathology (**Supplementary Fig. 10**). This highlights that EHT maturity is critical to modeling of pathological hypertrophy, which has also been observed previously with 2D-cultured iPSC-CMs^68^. Therefore, EHTs were cultured in MM for longer duration (i.e., 18 days) before treatment with agonists. We found that adrenergic agonist-treated EHTs had a 10-fold increase in gene expression of *NPPB* (encoding B-type natriuretic peptide, BNP) compared to untreated EHTs (**Fig. 5f**). This is particularly interesting as 2D iPSC-CMs at the same cell age as EHTs treated with the PE/ISO agonists failed to respond with no significant upregulation in either *NPPA* or *NPPB* (**Supplementary Fig. 11**). This highlighted that 3D EHTs provide a superior method to model pathological hypertrophy. Furthermore, we evaluated secreted BNP levels in the culture media by ELISA and observed that levels increased significantly both in the agonist-treated and elevated stiffness models compared to controls (**Fig. 5g, Supplementary Fig. 11**). The increase in secreted BNP in the chronic stiffness model despite no change in *NPPB* expression, suggests that these tissues had reached a new physiological baseline due to 18 days spent in stiff conditions. We expect that the initial spike in *NPPB* gene expression began immediately upon seeding the iPSC-CMs in stiff molds and then waned as secreted protein levels equilibrated to their maximum in the culture media. Although attempts to confirm cellular hypertrophy via imaging of cell boundaries were challenging, nuclear area from the cells in each condition was used as a proxy, since nucleus size has previously been shown to positively correlate with cell size^69^. This analysis showed that both treatment conditions led to a significant increase in nuclear area compared to controls (**Fig. 5h**). Western blot analysis also revealed increased activation of kinases linked to pathology, including the ERK1/2 and P38 MAP kinases (**Fig. 5i-j**). Activation of ERK and P38 signaling is well-established in human heart disease^7^. Interestingly, there were also some differences in pathological signaling between the two models as activation of Ca^2+^/calmodulin-dependent protein kinase II was only observed in the chronic stiffness-treated tissues (**Fig. 5i-j**).

Finally, we examined the effect of exposure to these pathological stressors on EHT function. Under electrical pacing, the relaxation time decreased significantly when EHTs were treated with agonists or exposed to overload from increased stiffness (**Fig. 5k**). Additionally, EHTs treated with agonists generated more contractile force when compared to untreated EHTs (**Fig. 5k**), an expected result with exposure to adrenergic agonism^70,71^. Measurement of contractile force in EHTs formed within stiff micropillar molds was not possible due to minimal stiff pillar deflection relative to the force generated (**Fig. 5k**). In addition to analysis of tissue contraction via bright field imaging, we performed analysis of calcium traces under spontaneous beating and electrical stimulation (1 Hz and 2 Hz). Calcium FWHM decreased for EHTs treated with adrenergic agonists or stiffness overload when paced at 2 Hz, whereas it increased for treated EHTs under unstimulated or spontaneous contractions (**Fig. 5l, Supplementary Fig. 11**). This accelerated calcium handling in response to pathological stressors matches previous reports that introduced increased resistance to EHT contraction via increased pillar stiffness^66^. Discrepancies can arise due to variability between iPSC lines, differences in cardiomyocyte population between replicates, and measurement techniques. The normalized contraction amplitude increased in the adrenergic agonist-treated EHTs as expected with chronic exposure to positive ionotropic drugs; however, no significant changes in normalized contraction amplitude were observed in EHTs exposed to stiffness overload (**Fig. 5l**). Furthermore, unlike the robust sarcomere assembly and expression in untreated EHTs, those exposed to stiffness overload displayed irregular sarcomere organization indicative of a disease phenotype (**Supplementary Fig. 11**). These collective findings indicate that, in addition to modeling molecular features of disease, these microtissues exhibit expected functional phenotypes in response to pathological stressors.

## OUTLOOK

While numerous *in vitro* cardiac models have been developed to study disease, their fabrication often requires multiple steps, which contributes to increased financial costs and limited adaptability due to logistical limitations. To address the limitations of scale-up and robustness, we used DLP-based 3D printing for one-step fabrication of hydrogel micropillar devices to form EHTs with high precision and efficiency (> 95%). Direct printing of hydrogels allowed for important mold properties such as high water content and tunable stiffness, which can be challenging with previously used elastomers. We first demonstrated the design flexibility afforded by DLP to fabricate EHTs with varying shapes (e.g., dog-bone, rings) and aspect ratio including scale-up to a 48-well format, which expands the applicability of the method to many EHT designs. Metabolic-driven maturation of 3D EHTs through use of a low glucose culture medium supplemented with fatty acids resulted in increased β-oxidation genes and improved sarcomere expression and alignment compared to 2D seeded iPSC-CMs, which illustrates the applicability of the EHTs to model cardiac tissue. Given the feasibility of fabrication and improved maturation, the 3D EHT approach was next applied for *in vitro* modeling of pathological cardiac hypertrophy (i.e., acute adrenergic agonism or chronic culture within stiff hydrogel molds) and treated EHTs exhibited increased levels of pathology-associated gene expression compared to untreated controls or treated 2D iPSC-CMs. Additionally, treated EHTs secreted BNP at levels similar to patients with heart failure and displayed activation of proteins involved in pathological remodeling, including ERK, P38, and CaMKII. This highlights robust induction of hypertrophic gene program in our 3D model compared to other methods, illustrating the potential use to model disease and to be used in approaches such as drug screening to identify new therapeutics.

This work opens up DLP printing as a facile and adoptable approach to fabricate microtissues to study a wide array of cardiac diseases. Future iterations of this platform can incorporate additional cardiac cell types (e.g., endothelial, immune cells, pericytes) to capture the effects of paracrine signaling on EHT performance. Further, the complexity of the 3D EHTs can be further enhanced through inclusion of vasculature, electrophysiological stimulation during culture, and sex-specificity compared to the human heart depending on the desired future application. Beyond the high speed of fabrication and ease of personalized design, DLP printing of hydrogel micropillar molds can also open avenues for integration of multi-functional materials (e.g., electrodes for stimulation during culture) in the future to enable further maturation to an adult phenotype and modeling of other cardiac diseases. Lastly, although this work focused on cardiac tissues and disease, the various technologies developed can be used in a vast array of tissues and diseases.

## METHODS

### Ethical Oversight

Animal experiments were approved by the University of Colorado Boulder Institutional Animal Care and Use Committee (IACUC). The University of Colorado Boulder animal facility is accredited by the Association for Assessment and Accreditation of Laboratory Animal Care (AAALAC). Non-failing human heart tissues were procured from donor hearts that failed due to age, size, or other incompatibility. Tissue collection, storage, and analysis was approved by the Colorado Multiple Institutional Review Board.

### Mouse Model of Pathological Cardiac Hypertrophy

Sixteen eight-week-old male C57Bl6/J mice were randomly assigned (eight per group) to receive either the β-adrenergic agonist isoproterenol (ISO) at 30 mg per kg bodyweight per day, or equal volume vehicle (1 µM L-ascorbic acid in sterile saline), to induce pathological cardiac remodeling. Continuous delivery was achieved by osmosis through subcutaneous implantation of a mini-osmotic pump containing ISO or vehicle posterior to the scapulae^72^. Seven days after pump implantation, the mice were humanely euthanized by cardiac explant under deep isoflurane-induced anesthesia. The explanted heart was rinsed with ice-cold saline, blotted dry, and then whole heart and left ventricle weights were measured. The sectioned cardiac chambers were snap frozen in liquid nitrogen and stored at -80°C.

### Protein Extraction

#### From iPSC-CMs and EHTs

Protein lysates were collected from 2D iPSC-CMs by adding 200 µl RIPA buffer (Invitrogen) to each well, scraping, and collecting in a microcentrifuge tube. EHTs were harvested by collecting microtissues from microwells and placing into 100 µl of RIPA buffer in a microcentrifuge tube. Lysates from both conditions were incubated rotating at 4°C for three hours. Samples were boiled at 95°C for 10 minutes and then centrifuged at 13,000 ×g for seven minutes. Supernatants were collected and stored at -80°C and protein concentration was determined by BCA Assay (Pierce).

#### From Mouse Cardiac Tissue

Mouse left ventricular tissue (∼50 mg) was pulverized while frozen using a BioPulverizer (Bio Spec) and then added to Eppendorf tubes containing RIPA buffer (Invitrogen) supplemented with protease and phosphatase inhibitors (Halt). Lysis was achieved by overnight incubation at 4°C with end-over-end agitation. The samples were then centrifuged at 12,000 ×g for 10 minutes at 4°C and the supernatants collected for downstream analyses. Protein concentration was determined by BCA Assay (Pierce).

### RNA Extraction

#### From iPSC-CMs and EHTs

RNA from 2D iPSC-CMs was collected by adding 300 µl Buffer RLT (Qiagen) to wells, scraping, and then collecting into microcentrifuge tube. For EHTs, microtissues from each condition were extracted from microwells and added to 300 µl of Buffer RLT. All samples were incubated rotating at 4°C for three hours. The samples were then centrifuged at 12,000 ×g for five minutes. Supernatant was combined with 70% ethanol and added onto column from RNeasy Mini Kit (Qiagen). Washes and RNA elution were performed according to the manufacturer provided protocol. RNA concentration was determined by Nanodrop.

#### From Cardiac Tissue

Left ventricular tissue (∼40 mg) was added to 1 mL of Tri Reagent and homogenized with a mechanical homogenizer (IKA Works, T10 Basic S1) at 20,000 RPM. After incubating for five minutes at room temperature, 200 µL chloroform was added, the tubes shaken vigorously for ∼15 seconds, and then incubated at room temperature for 15 minutes. The samples were then centrifuged at 12,000 ×g for 15 minutes at 4°C and the upper aqueous layer (∼500 µL) was collected into a clean Eppendorf tube containing 500 µL of isopropanol. The samples were briefly vortexed to mix and then incubated at -20°C to improve RNA precipitation. The RNA was then pelleted by centrifugation at 12,000 RCF for 10 minutes at 4°C. The supernatant was discarded, and the RNA pellet was washed twice by resuspension in ice-cold 75% ethanol followed by centrifugation. After discarding the second ethanol wash, the RNA was dried for ∼10 minutes by uncapping the tubes. RNA was then solubilized in milli-Q water. RNA concentration and quality were assessed by Nanodrop.

### iPSC Cardiomyocyte Differentiation and Culture

WTC11 iPSC line was obtained from the Coriell Institute and differentiated into cardiomyocytes using a previously well-established protocol with minor modifications^73^. In brief, iPSCs were cultured in mTSER+ media until 70% confluency was reached and cells were split onto 12-well plates. Once iPSCs reached between 50-70% confluency on a 12-well plate, the differentiation protocol was initiated. For differentiation Day 0 to Day 6 the base media was RPMI 1640 medium supplemented with ascorbic acid (5 mg mL^-1^) and BSA (40 mg mL^-1^). On Day 0, the cells were treated with the Wnt activator Chiron 99021 (Caymen) at 5 µM for 48 hours. The cells were washed once with PBS and then the Wnt inhibitor C59 added at 2 µM for 48 hours. After these Wnt modulation steps, the media was changed to base medium for 48 hours. On Day 6 of differentiation, the media was changed to RPMI 1640 containing B27 (1×) and penicillin–streptomycin (1% v/v). Media was changed every two days until cells were reseeded onto EHTs between days 10 and 12.

### 3D Printing and Mechanical Characterization of Hydrogel Micropillar Molds

All CAD models for hydrogel micropillar molds used in the study were designed in Fusion 360 (Autodesk, USA). Poly(ethylene glycol) diacrylate (PEGDA, molecular weight = 700 Da, 10 wt.%), lithium phenyl-2,4,6-trimethylbenzoylphosphinate (LAP, 0.5 wt.%), tartrazine (TTz, 2 mM) and Dulbecco’s phosphate buffered saline (DPBS) were used as macromer, photoinitiator, photoabsorber, and solvent, respectively, to formulate the resin for printing of soft hydrogel micropillar molds. 3D printing of hydrogel molds was conducted on a Lumen Alpha DLP printer (35 μm xy pixel resolution, Volumetric Inc., USA) as described previously^74^. Briefly, PEGDA resin was dispensed into the poly(dimethyl siloxane) PDMS vat, and the layers were crosslinked sequentially with light exposure (7.5 s, 20 mW cm^-2^, 100 μm layer thickness). The base layers (200 μm layer thickness) were irradiated at 15 s to facilitate attachment of the mold to the build platform. After printing was complete, 3D printed hydrogels were removed from the build platform and immersed in DPBS on a shaker to remove unreacted monomer, photoinitiator, and photoabsorber. After repeated washing and rinsing, the molds were sterilized in 70% ethanol for 30 minutes, followed by sterilization under UV germicidal lamp for another 30 minutes, and stored in sterile PBS at 4 °C until use. 3D printing of hydrogel micropillar molds was undertaken on a Sonic Mini 8K S LCD printer (22 μm xy pixel resolution, Phrozen Inc., Taiwan) for comparison. PEGDA resin formulation as mentioned earlier was dispensed directly into the vat and printing was subsequently conducted with light exposure (55 s, 2.35 mW cm^-2^, 100 μm layer thickness). Stiff micropillar molds were 3D printed using PEGDA Start Photoink™ (Cellink Inc.) following manufacturer’s protocol.

In situ photo-rheological (oscillatory shear time sweeps) characterization was performed on a Discovery HR20 rheometer (TA Instruments) (equipped with 20 mm diameter parallel plate geometry, 100 µm gap, 25 °C) to assess time to reach gelation (defined by the crossover between storage, *G*′ and loss modulus, *G*″) in the presence of visible light (20 mW cm^-2^, Exfo Omnicure Vis S1000 lamp with 400 nm to 500 nm bandpass filter). Unconfined compression testing (Q800 DMA, TA Instruments, force ramp = 0.5 N min^-1^) was performed on 3D printed hydrogel discs (5 mm diameter, 1.5 mm thickness). Compressive moduli were determined from the linear elastic region (10-20% strain) of the stress–strain curves. Uniaxial tensile testing until failure (RSA-G2 DMA, TA Instruments, stretch rate = 0.05 s^-1^) was performed on 3D printed rectangular strips (15 × 5 × 1.5 mm^3^). Tensile moduli (slope from 5–15% strain) were determined from the stress–stretch curves using TA Instrument Analysis software. All samples were swollen to equilibrium before testing. Polymer content of hydrogels in the equilibrium swollen state was determined using equation 1 as reported previously^75^.

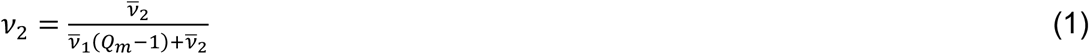

Where, mass swelling ratio (*Q*_*m*_) was calculated as the ratio of the swollen gel mass to that of dry polymer after lyophilization. *⊽*_1_ is the specific volume of water (*⊽*_1_ = 1 mL g^-1^) and *⊽*_*2*_ is the specific volume of poly(ethylene glycol) diacrylate (*⊽*_2_ = 0.87 mL g^-1^)

Computational modeling of micropillar bending was performed on Autodesk Fusion 360 using a static stress simulation. Briefly, the displacement of hydrogel micropillar was determined under varying horizontal force (1 to 500 μN). To mimic the force applied by the EHTs, simulated force was applied perpendicular to the pillar in the form of a planar rectangular section (0.2 mm height, 0.3 mm width, placed 0.33 mm from the bottom of the well). Custom material settings were added into the physical material library for the hydrogel micropillar CAD model. The material ’s mechanical attributes including Young’s modulus (= 51 kPa), Poisson’s ratio (= 0.36), shear modulus (= 16 kPa), density (= 1.15 g cm^-3^), damping coefficient (= 0), and tensile strength (= 17 kPa) were taken into consideration within the model. Stiffness of the micropillars was determined as the ratio of the applied force to measured maximum deflection.

### Fabrication and Treatment of EHTs

Hydrogel micropillar molds were aspirated to remove any remaining liquid from the wells. Acid-solubilized type I bovine telocollagen (Advanced BioMatrix) was mixed with prechilled 10× DPBS, sterile distilled water, and neutralized with 0.1 N NaOH to achieve a final concentration of 1 mg mL^-1^. The solution was supplemented with Matrigel basement membrane matrix (10%, growth factor reduced, Corning) and kept under ice-cold conditions. iPSC-CMs were harvested with trypsin-EDTA (0.25%, Gibco) at 37 °C for 7 minutes. Recovery media containing fetal bovine serum (FBS) was added to wells to neutralize the trypsin, cells were gently collected and pelleted via centrifugation (300×g for 3 minutes). Supernatant was aspirated and cells were resuspended in neutralization media. iPSC-CM cell suspension (cell age: day 12 to 14, density: 10 million cells mL^-1^) was mixed with collagen precursor, 5 to 7 μL of the resultant solution was transferred to micropillar molds, and the solution was incubated at 37 °C in a 5% CO2 incubator for 15 min to undergo gelation. After incubation, the hydrogel micropillar molds containing cell-laden gels were immersed in recovery medium comprising RPMI 1640 (Gibco) supplemented with B-27 (1×, Thermo Scientific), FBS (20% v/v), and penicillin–streptomycin (1% v/v). To visualize the collagen matrix in the EHTs, FITC-conjugated collagen (Sigma Aldrich, C4361, 0.08 mg mL^-1^) was mixed with nonlabelled collagen (0.92 mg mL^-1^). The resultant collagen mixture was neutralized and mixed with cells as described earlier^74^. To form fibrin-based EHTs, fibrinogen (from human plasma, 4 mg mL^-1^) was mixed with ice-cold solution of DPBS, Geltrex, and iPSC-CMs. The resulting solution was mixed with thrombin (0.4 U mL^-1^) immediately before seeding, placed in an incubator (37 °C, 5% CO2) for 90 minutes to allow crosslinking, and then incubated in media containing aprotinin (0.033 mg mL^-1^). The day of seeding cell-laden gels within micropillar molds is defined as day 0. Day 2 after seeding, the recovery medium was replaced with growth medium consisting of RPMI 1640 with B-27 (1×) and penicillin–streptomycin (1% v/v) with regular media changes every other day for the first 10 days. After growth phase at day 10, the media was replaced with maturation media consisting of glucose free RPMI supplemented with low glucose (mM), palmitic acid (PA, 50 𝜇M), oleic acid (OA, 100 𝜇M), galactose (10 mM), B27 (1×), and penicillin–streptomycin (1% v/v). EHTs were cultured in maturation media until day 28 (∼cell age: day 40) before starting any treatment. To induce pathological cardiac hypertrophy via soluble factors, EHTs were treated with a cocktail of adrenergic agonists: phenylephrine (PE, 100 𝜇M) and isoproterenol (ISO, 20 𝜇M). To induce pathological cardiac hypertrophy via stiffness overload, EHTs were formed within stiff hydrogel micropillar molds and cultured for the entire duration without agonists. EHTs were pooled within each group and harvested one day after treatment (day 29) to isolate RNA or protein, respectively. Analysis of EHT function (contractility and calcium handling) was performed seven days after treatment (day 35). Conditioned media in each condition was collected at days 1, 3, 7 post treatment for analysis of secreted factors.

### Immunostaining of EHTs

EHTs at relevant time points were fixed with neutral buffered formalin (10% v/v) for 30 minutes, permeabilized with Triton X-100 (0.1% v/v), blocked with BSA (5 wt.%), and stained with fluorescein-conjugated phalloidin (1:500) and Hoechst 33342 (1:1000) to visualize F-actin and nucleus, respectively. Microtissues were stained with primary antibodies for cardiac troponin-T (1:200), vimentin (1:200), and sarcomeric α-actinin (1:200) to visualize the relative proportion of cardiomyocytes to non-myocytes and sarcomere, respectively (**Supplementary Table 1**). Z-stack fluorescence images were acquired on a confocal microscope (Nikon Eclipse Ti2 with AXR scanner) equipped with a 2× or 20× objective lens. Images were analysed using ImageJ (NIH) and linear adjustment of brightness/contrast was applied to the entire image. Sarcomere length was measured for cells stained with sarcomeric α-actinin by measuring the distance between intensity peaks along the long axis (i.e., distance between the center of adjacent z-discs). The dominant direction of alignment and distribution of the sarcomere as well as the collagen fibres within EHTs were analysed using the Directionality and OrientationJ plugins, respectively in Fiji (ImageJ). To assess viability, EHTs were incubated with calcein AM and ethidium homodimer-1 at 37 °C for 30 minutes per manufacturer’s protocol (Live/Dead™ Cytotoxicity Kit, Thermo Fisher Scientific). Fluorescence images were auto-threshold using Otsu filter to isolate total nuclei (Hoechst 33342) and dead cells (ethidium homodimer-1) analysed using ImageJ (NIH). The number of nuclei stained with ethidium homodimer-1 relative to the total number of nuclei were used to estimate the proportion of viable cells within EHTs. To evaluate fatty acid uptake, live EHTs in growth or maturation media were incubated in DPBS with BODIPY FL-C16 (1 μM, Thermo Scientific) for 45 minutes at 37 °C, washed 3× with DPBS to remove staining solution, and imaged on the confocal microscope using consistent settings (laser power and gain). Fluorescence images were analysed for mean gray area value using ImageJ to quantify the intensity of stain across experimental groups.

### Transmission Electron Microscopy of EHTs

EHTs were fixed were fixed using 3% paraformaldehyde, 0.35% glutaraldehyde, in 0.1 M sodium cacodylate buffer (pH 7.4, Electron Microscopy Sciences) for 45 min, then washed with 0.1 M sodium cacodylate buffer pH 7.4, and stored at 4 °C overnight. EHTs were removed from micropillar molds followed by post-fixing with osmium tetroxide (1% v/v) in 0.1 M sodium cacodylate for 1 hour, washed, and en-bloc stained for 30 min with aqueous uranyl acetate (2% v/v). Samples were serially dehydrated in ethanol over 4 hours, exchanged into anhydrous acetone, followed by Epon/Araldite infiltration over 5 days at room temperature, and then polymerized for 48 hours at 60 ºC. Thin sections (90-100 nm) were cut using a Leica UCT ultramicrotome. Serial sections were collected on Formvar-coated, copper slot grids, post-stained using aqueous uranyl acetate (2% v/v) followed by Reynold’s lead citrate and imaged using an FEI Tecnai T12 Spirit TEM, operating at 100 kV.

### Contractility and calcium handling of EHTs

Live cell, bright field videos of tissue contractility under electrical stimulation (1 to 3 Hz, 10 V cm^-1^, 37 °C with 5% CO_2_) were recorded. IonOptix electrodes with Myopacer pacing setup in a 6-well glass bottom plate was used to electrically stimulate the EHTs while imaging. The tissues were subjected to each stimulation frequency for 5 min before recording any measurements. The contraction recordings were processed through an open-source MuscleMotion software to determine the contraction amplitude, time to peak, and relaxation time^76^. Relaxation time was defined as the time required for EHT to relax from the peak contraction amplitude. Analysis of pillar deflection was conducted in a semi-automated fashion using Beatprofiler^77^. Briefly, pillars were manually labelled in ImageJ to identify the coordinates, and the data (microns per pixel, baseline unloaded distance between pillars) was input within the algorithm. The product of the output values for maximum pillar deflection and micropillar stiffness coefficient was used to calculate the contractile force. Calcium imaging and analysis were performed on EHTs at relevant time points. EHTs were treated with intracellular calcium indicator (Cal 520, AAT Bioquest, 4 µM) for 90 min (at 37 °C in a 5% CO2 incubator) in maintenance (growth or maturation) media supplemented with Pluronic (0.04%) to enhance cellular uptake. Following calcium loading, the staining solution was replaced with fresh maintenance media and videos acquired on Nikon Ti2 Eclipse with AXR scanning confocal microscope (2× objective, 105 Hz frame rate). The temporal change in fluorescence intensity was regarded as the calcium transient trace and used for all measurements. Calcium activation maps were produced by quantitative analysis of the captured videos using open-source software package ElectroMap as described previously^78^. Analysis of full width half maximum (FWHM) and calcium amplitude was performed through an automated software – Beatprofiler as described previously^77^. Calcium transient amplitude was defined as the area under the peak maxima relative to the baseline.

### Western Blotting

After determination of protein concentration by BCA Assay, 10 µg of protein from EHTs/iPSC-CMs or 20 µg of protein from the mouse LV was added to equal volume SDS loading buffer (Invitrogen) containing DTT reducing buffer (Bolt) and boiled for five minutes. Samples were then loaded onto 4-12% Bis-Tris gradient gels and protein separated by electrophoresis at 120 V. Proteins were then transferred to nitrocellulose membranes and Revert total protein stain (LI-COR) used to assess quality of transfer and serve as a loading control for downstream quantification. The membranes were then blocked with 5% milk for one hour at room temperature. Primary antibodies (**Supplementary Table 1**) were then added in 5% BSA and incubated overnight at 4°C with agitation. The membranes were washed with Tween tris buffered saline (TBS-T) and then secondary antibodies in 5% milk added for one hour at room temperature. The membranes were again washed with TBS-T and then imaged using an Azure c600 imager. Protein densitometry was measured using LI-COR Image Studio or Fiji (ImageJ).

### Enzyme-linked Immunosorbent Assay (ELISA)

The BNP Human Elisa Kit (abcam) was used to determine secreted levels of BNP in the conditioned media from control and agonist-treated EHTs following the manufacturer’s protocol. In brief, media was collected from appropriate wells and frozen at -80°C until ready for use. Standards, reagents, and samples were prepared according to manufacturer’s protocol. Wells were incubated with constant shaking overnight at 4°C. After following manufacturer’s protocol to prepare the sample plate, the wells were read at 450 nm on plate reader.

### In-Gel Digestion Mass Spectrometry

Approximately 20 µg of protein was boiled in SDS loading buffer containing Bolt reducing buffer and protease/phosphatase inhibitors and loaded onto a 4-12% Bis-Tris gradient gel. After separation of proteins by electrophoresis, the gel was fixed in 50% methanol/10% acetic acid for 15 minutes and then stained with Coomassie Brilliant Blue for 20 minutes. The gel was then washed overnight in 10% methanol/7.5% acetic acid at room temperature. When sufficiently de-stained, the protein band at ∼220 kDa (representing myosin heavy chain) was excised from the gel with a scalpel, diced into eight small pieces, and added to a clean Eppendorf tube. Protein gel bands were excised and destained with 50mM ammonium bicarbonate/50% (v/v) acetonitrile, dehydrated with 100% acetonitrile, reduced and alkylated with 10 mM TCEP and 40 mM 2-chloroacetamide in 50 mM ammonium bicarbonate at 70°C, then digested overnight with trypsin (Promega) overnight all at ambient. Tryptic peptides were extracted with 25% (v/v) acetonitrile/0.1% (v/v) trifluoroacetic acid, and 100% acetonitrile, then cleaned up using rpC18. Cleaned-up peptide were then dried in a Speedvac vacuum centrifuge and stored at -20°C until analysis. Samples were suspended in 3% (v/v) acetonitrile/0.1% (v/v) trifluoroacetic acid and approximately 1 picomole tryptic peptides were directly injected onto a rpC18 1.7 µm, 130 Å, 75 µm X 250 mm BEH M-class column (Waters), using a Waters M-class UPLC. Peptides were eluted at 300 nL min^-1^ using a gradient from 2% to 20% acetonitrile in 20 minutes and detected by an Orbitrap Fusion Tribrid mass spectrometer (Thermo Scientific) running Tune 3.3.2782.34. Precursor mass spectra (MS1) were acquired at a resolution of 120,000 from 350 to 1500 m/z with an automatic gain control (AGC) standard target and an auto maximum injection time. Precursor peptide ion isolation width for MS2 fragment scans was 1.6 m/z with a 3 second cycle time. All MS2 spectra were acquired at a resolution of 15,000 with higher energy collision dissociation (HCD) at 30% normalized collision energy. The AGC target was standard and the maximum injection time was auto. Rawfiles were searched against the Uniprot Human database UP000005640 (downloaded 08/26/2022) using Maxquant version 2.0.3.0 with cysteine carbamidomethylation as a fixed modification. Methionine oxidation and protein N-terminal acetylation were searched as variable modifications. All peptides and proteins were thresholded at a 1% false discovery rate (FDR).

### Complementary DNA Synthesis and Quantitative RT-PCR

RNA from 2D iPSC-CMs and EHTs were used to make cDNA using SuperScript III reverse transcriptase (Invitrogen). In brief, 150 ng-500 ng of mRNA was used for cDNA synthesis with random hexamer primers and SuperScript III reverse transcriptase. For qPCR, 5 ng cDNA was combined with 500 nM each forward and reverse primers and SYBR Green PCR Master Mix (Thermo). Genes of interest were normalized to housekeeping gene GAPDH. Data was analysed using the 2^Λ1Λ1CT method. The sequences of the primers used in the qPCR analyses are available in **Supplementary Table 2**.

### RNA Sequencing

Total RNA samples were sent to Novogene Corporation (Sacramento, CA) for poly-A enrichment, cDNA library preparation, and mRNA sequencing. mRNA sequencing was performed on an Illumina Novaseq 6000 with 150 base-pair reads and a read depth of 50 million paired-end reads. For analysis, Nextflow version 21.10.6 was used to run the nf-core/rnaseq pipeline version 3.9.0^79,80^. Raw gene counts were generated by salmon version 1.5.2^81^. Gene count normalization and differential gene expression analysis was conducted with the R (version 4.2) package DESeq2 (version 1.36.0)^82^. Prefiltering was conducted to remove genes that did not have at least 10 counts for 2 samples. Pairwise comparisons between 2D and EHT samples were conducted and the apeglm shrinkage estimator was used to produce shrunken log fold changes for visualization^83^. Differentially expressed genes were defined as those with an adjusted p-value < 0.05 and an absolute shrunken fold change greater than 1.5. Relative proportions of isoforms were calculated using transcript abundance values generated by salmon.

### Statistical Analysis

Comparisons of two groups were performed using an unpaired Student’s two-tailed t-test. For comparisons of three or more groups with one categorical variable, a one-way analysis of variance (ANOVA) was used. When a significant interaction was identified, Tukey’s post-hoc test for multiple independent pairwise comparisons was used. These analyses were performed using GraphPad Prism version 10. A p value of < 0.05 was considered statistically significant.

## DATA AVAILABILITY

The main data supporting the results in this study are available within the paper and its Supplementary Information. Source data for the figures will be provided with this paper.

## ACKNOWLEDGEMENTS

Electron microscopy was performed at the University of Colorado, Boulder EM Services Core Facility in the MCDB Department and the authors are grateful for technical assistance of facility staff (Sarah Zimmerman and Garry Morgan). The authors are grateful to Dr. Thomas Kolibaba and Dr. Jason Killgore for assistance with measurement of spectral properties; Dr. Massimo Buvoli, Dr. Kristen Bjorkman, Dr. Yuxiao Tan, Dr. Matthew Davidson, Dr. Manuela Garay-Sarmiento, and Bruce Kirkpatrick for helpful discussions.

## AUTHOR CONTRIBUTIONS

A.P.D., M.A.J., T.G.M., J.A.B., and L.A.L. conceptualized and designed the project. A.P.D., M.A.J., T.G.M. performed experiments and conducted data analysis. G.J.R-R. and B.M-Z. provided experimental assistance with imaging and analysis of tissue contractility. D.R.H. conducted analysis of RNA-Seq data. M.C.O assisted with experiments and data analysis pertaining to changing cell density and matrix type. C.O.C. conducted optimization of mold design and Q.M. assisted with transmission electron microscopy of tissues. H.V-A. and C.C.E assisted with mass spectrometry experiments. H.M.Z. and D.N.G performed computational analysis of pillar deflection. J.A.B. and L.A.L. supervised the entire study and acquired funding. A.P.D., M.A.J., T.G.M. prepared the initial draft of the manuscript with inputs from J.A.B. and L.A.L. All authors contributed to reviewing and editing of the final manuscript.

## REFERENCES

1. Ingber, D. E. Human organs-on-chips for disease modelling, drug development and personalized medicine. Nature Reviews Genetics 2022 23:8 23, 467–491 (2022).

2. Zushin, P. J. H., Mukherjee, S. & Wu, J. C. FDA Modernization Act 2.0: transitioning beyond animal models with human cells, organoids, and AI/ML-based approaches. J Clin Invest 133, (2023).

3. Sharma, A., Sances, S., Workman, M. J. & Svendsen, C. N. Multi-lineage Human iPSC-Derived Platforms for Disease Modeling and Drug Discovery. Cell Stem Cell 26, 309–329 (2020).

4. Fonseca, A. C. et al. Emulating Human Tissues and Organs: A Bioprinting Perspective Toward Personalized Medicine. Chem Rev 120, 11128–11174 (2020).

5. Ronaldson-Bouchard, K. & Vunjak-Novakovic, G. Organs-on-a-Chip: A Fast Track for Engineered Human Tissues in Drug Development. Cell Stem Cell 22, 310–324 (2018).

6. Moroni, L. et al. Biofabrication strategies for 3D in vitro models and regenerative medicine. Nature Reviews Materials 2018 3:5 3, 21–37 (2018).

7. Martin, T. G., Juarros, M. A. & Leinwand, L. A. Regression of cardiac hypertrophy in health and disease: mechanisms and therapeutic potential. Nature Reviews Cardiology 2023 20:5 20, 347–363 (2023).

8. Thavandiran, N. et al. Design and formulation of functional pluripotent stem cell-derived cardiac microtissues. Proc Natl Acad Sci U S A 110, E4698–E4707 (2013).

9. Cho, S., Lee, C., Skylar-Scott, M. A., Heilshorn, S. C. & Wu, J. C. Reconstructing the heart using iPSCs: Engineering strategies and applications. J Mol Cell Cardiol 157, 56–65 (2021).

10. Kim, H., Kamm, R. D., Vunjak-Novakovic, G. & Wu, J. C. Progress in multicellular human cardiac organoids for clinical applications. Cell Stem Cell 29, 503–514 (2022).

11. Legant, W. R. et al. Microfabricated tissue gauges to measure and manipulate forces from 3D microtissues. Proc Natl Acad Sci U S A 106, 10097–10102 (2009).

12. Nunes, S. S. et al. Biowire: a platform for maturation of human pluripotent stem cell–derived cardiomyocytes. Nature Methods 2013 10:8 10, 781–787 (2013).

13. Ronaldson-Bouchard, K. et al. Advanced maturation of human cardiac tissue grown from pluripotent stem cells. Nature 2018 556:7700 556, 239–243 (2018).

14. Schwan, J. et al. Anisotropic engineered heart tissue made from laser-cut decellularized myocardium. Scientific Reports 2016 6:1 6, 1–12 (2016).

15. Skylar-Scott, M. A. et al. Biomanufacturing of organ-specific tissues with high cellular density and embedded vascular channels. Sci Adv 5, (2019).

16. Lind, J. U. et al. Instrumented cardiac microphysiological devices via multimaterial three-dimensional printing. Nature Materials 2016 16:3 16, 303–308 (2016).

17. Liu, J. et al. Direct 3D bioprinting of cardiac micro-tissues mimicking native myocardium. Biomaterials 256, 120204 (2020).

18. Daly, A. C., Davidson, M. D. & Burdick, J. A. 3D bioprinting of high cell-density heterogeneous tissue models through spheroid fusion within self-healing hydrogels. Nature Communications 2021 12:1 12, 1–13 (2021).

19. Hamidzada, H. et al. Primitive macrophages induce sarcomeric maturation and functional enhancement of developing human cardiac microtissues via efferocytic pathways. Nature Cardiovascular Research 2024 3:5 3, 567–593 (2024).

20. Landau, S. et al. Primitive macrophages enable long-term vascularization of human heart-on-a-chip platforms. Cell Stem Cell 31, 1222-1238.e10 (2024).

21. Michas, C. et al. Engineering a living cardiac pump on a chip using high-precision fabrication. Sci Adv 8, 3791 (2022).

22. Mathur, A. et al. Human iPSC-based Cardiac Microphysiological System For Drug Screening Applications. Scientific Reports 2015 5:1 5, 1–7 (2015).

23. Bannerman, D. et al. Heart-on-a-Chip Model of Epicardial–Myocardial Interaction in Ischemia Reperfusion Injury. Adv Healthc Mater 13, 2302642 (2024).

24. Mannhardt, I. et al. Comparison of 10 Control hPSC Lines for Drug Screening in an Engineered Heart Tissue Format. Stem Cell Reports 15, 983–998 (2020).

25. Wang, G. et al. Modeling the mitochondrial cardiomyopathy of Barth syndrome with induced pluripotent stem cell and heart-on-chip technologies. Nature Medicine 2014 20:6 20, 616–623 (2014).

26. Tu, C. et al. Tachycardia-induced metabolic rewiring as a driver of contractile dysfunction. Nature Biomedical Engineering 2023 8:4 8, 479–494 (2023).

27. Cho, S., Discher, D. E., Leong, K. W., Vunjak-Novakovic, G. & Wu, J. C. Challenges and opportunities for the next generation of cardiovascular tissue engineering. Nature Methods 2022 19:9 19, 1064–1071 (2022).

28. Ogle, B. M. et al. Distilling complexity to advance cardiac tissue engineering. Sci Transl Med 8, (2016).

29. Huebsch, N. et al. Miniaturized iPS-Cell-Derived Cardiac Muscles for Physiologically Relevant Drug Response Analyses. Scientific Reports 2016 6:1 6, 1–12 (2016).

30. Breckwoldt, K. et al. Differentiation of cardiomyocytes and generation of human engineered heart tissue. Nature Protocols 2017 12:6 12, 1177–1197 (2017).

31. Simmons, D. W., Schuftan, D. R., Ramahdita, G. & Huebsch, N. Hydrogel-Assisted Double Molding Enables Rapid Replication of Stereolithographic 3D Prints for Engineered Tissue Design. ACS Appl Mater Interfaces 15, 25313–25323 (2023).

32. Zhao, Y. et al. A Multimaterial Microphysiological Platform Enabled by Rapid Casting of Elastic Microwires. Adv Healthc Mater 8, 1801187 (2019).

33. Tamargo, M. A. et al. MilliPillar: A Platform for the Generation and Real-Time Assessment of Human Engineered Cardiac Tissues. ACS Biomater Sci Eng 7, 5215–5229 (2021).

34. Leung, C. M. et al. A guide to the organ-on-a-chip. Nature Reviews Methods Primers 2022 2:1 2, 1–29 (2022).

35. Campbell, S. B. et al. Beyond Polydimethylsiloxane: Alternative Materials for Fabrication of Organ-on-a-Chip Devices and Microphysiological Systems. ACS Biomater Sci Eng 7, 2880–2899 (2021).

36. Grigoryan, B. et al. Multivascular networks and functional intravascular topologies within biocompatible hydrogels. Science (1979) 364, 458–464 (2019).

37. Dhand, A. P. et al. Additive manufacturing of highly entangled polymer networks. Science (1979) 385, 566–572 (2024).

38. Levato, R. et al. Light-based vat-polymerization bioprinting. Nature Reviews Methods Primers 2023 3:1 3, 1–19 (2023).

39. Lim, K. S. et al. Fundamentals and Applications of Photo-Cross-Linking in Bioprinting. Chem Rev 120, 10662–10694 (2020).

40. Howard, C. M. & Baudino, T. A. Dynamic cell–cell and cell–ECM interactions in the heart. J Mol Cell Cardiol 70, 19–26 (2014).

41. Boudou, T. et al. A Microfabricated Platform to Measure and Manipulate the Mechanics of Engineered Cardiac Microtissues. https://home.liebertpub.com/tea 18, 910–919 (2011).

42. Zhao, Y. et al. A Platform for Generation of Chamber-Specific Cardiac Tissues and Disease Modeling. Cell 176, 913-927.e18 (2019).

43. Goldfracht, I. et al. Generating ring-shaped engineered heart tissues from ventricular and atrial human pluripotent stem cell-derived cardiomyocytes. Nature Communications 2020 11:1 11, 1–15 (2020).

44. Bose, P., Eyckmans, J., Nguyen, T. D., Chen, C. S. & Reich, D. H. Effects of Geometry on the Mechanics and Alignment of Three-Dimensional Engineered Microtissues. ACS Biomater Sci Eng 5, 3843–3855 (2019).

45. Eder, A., Vollert, I., Hansen, A. & Eschenhagen, T. Human engineered heart tissue as a model system for drug testing. Adv Drug Deliv Rev 96, 214–224 (2016).

46. Yang, X., Pabon, L. & Murry, C. E. Engineering adolescence: Maturation of human pluripotent stem cell-derived cardiomyocytes. Circ Res 114, 511–523 (2014).

47. Black, L. D., Meyers, J. D., Weinbaum, J. S., Shvelidze, Y. A. & Tranquillo, R. T. Cell-Induced Alignment Augments Twitch Force in Fibrin Gel-Based Engineered Myocardium via Gap Junction Modification. Tissue Eng Part A 15, 3099–3108 (2009).

48. Feric, N. T. & Radisic, M. Maturing human pluripotent stem cell-derived cardiomyocytes in human engineered cardiac tissues. Adv Drug Deliv Rev 96, 110–134 (2016).

49. Schaaf, S. et al. Human Engineered Heart Tissue as a Versatile Tool in Basic Research and Preclinical Toxicology. PLoS One 6, e26397 (2011).

50. Turnbull, I. C. et al. Advancing functional engineered cardiac tissues toward a preclinical model of human myocardium. The FASEB Journal 28, 644–654 (2014).

51. Karbassi, E. et al. Cardiomyocyte maturation: advances in knowledge and implications for regenerative medicine. Nature Reviews Cardiology 2020 17:6 17, 341–359 (2020).

52. Knight, W. E., Cao, Y., Dillon, P. & Song, K. A simple protocol to produce mature human-induced pluripotent stem cell-derived cardiomyocytes. STAR Protoc 2, 100912 (2021).

53. Knight, W. E. et al. Maturation of Pluripotent Stem Cell-Derived Cardiomyocytes Enables Modeling of Human Hypertrophic Cardiomyopathy. Stem Cell Reports 16, 519–533 (2021).

54. Huebsch, N. et al. Metabolically driven maturation of human-induced-pluripotent-stem-cell-derived cardiac microtissues on microfluidic chips. Nature Biomedical Engineering 2022 6:4 6, 372–388 (2022).

55. Reiser, P. J., Portman, M. A., Ning, X. H. & Moravec, C. S. Human cardiac myosin heavy chain isoforms in fetal and failing adult atria and ventricles. Am J Physiol Heart Circ Physiol 280, (2001).

56. Martin, T. G. & Kirk, J. A. Under construction: The dynamic assembly, maintenance, and degradation of the cardiac sarcomere. J Mol Cell Cardiol 148, 89–102 (2020).

57. Crocini, C. & Gotthardt, M. Cardiac sarcomere mechanics in health and disease. Biophysical Reviews 2021 13:5 13, 637–652 (2021).

58. Litviňuková, M. et al. Cells of the adult human heart. Nature 2020 588:7838 588, 466–472 (2020).

59. Wright, C. J., Smith, C. W. J. & Jiggins, C. D. Alternative splicing as a source of phenotypic diversity. Nature Reviews Genetics 2022 23:11 23, 697–710 (2022).

60. Huang, J. et al. Regulation of Postnatal Cardiomyocyte Maturation by an RNA Splicing Regulator RBFox1. Circulation 148, 1263–1266 (2023).

61. Duran, J., Nickel, L., Estrada, M., Backs, J. & van den Hoogenhof, M. M. G. CaMKIIδ Splice Variants in the Healthy and Diseased Heart. Front Cell Dev Biol 9, 644630 (2021).

62. Kong, S. W. et al. Heart Failure-Associated Changes in RNA Splicing of Sarcomere Genes. Circ Cardiovasc Genet 3, 138–146 (2010).

63. Cejas, R. B., Tamaño-Blanco, M., Fontecha, J. E. & Blanco, J. G. Impact of DYRK1A Expression on TNNT2 Splicing and Daunorubicin Toxicity in Human iPSC-Derived Cardiomyocytes. Cardiovasc Toxicol 22, 701–712 (2022).

64. Rodriguez, M. L., Werner, T. R., Becker, B., Eschenhagen, T. & Hirt, M. N. Magnetics-Based Approach for Fine-Tuning Afterload in Engineered Heart Tissues. ACS Biomater Sci Eng 5, 3663–3675 (2019).

65. Nakamura, M. & Sadoshima, J. Mechanisms of physiological and pathological cardiac hypertrophy. Nature Reviews Cardiology 2018 15:7 15, 387–407 (2018).

66. Leonard, A. et al. Afterload promotes maturation of human induced pluripotent stem cell derived cardiomyocytes in engineered heart tissues. J Mol Cell Cardiol 118, 147–158 (2018).

67. Hirt, M. N. et al. Increased afterload induces pathological cardiac hypertrophy: A new in vitro model. Basic Res Cardiol 107, 1–16 (2012).

68. Johansson, M. et al. Cardiac hypertrophy in a dish: A human stem cell based model. Biol Open 9, (2020).

69. Wu, Y., Pegoraro, A. F., Weitz, D. A., Janmey, P. & Sun, S. X. The correlation between cell and nucleus size is explained by an eukaryotic cell growth model. PLoS Comput Biol 18, e1009400 (2022).

70. Ergir, E. et al. Generation and maturation of human iPSC-derived 3D organotypic cardiac microtissues in long-term culture. Scientific Reports 2022 12:1 12, 1–21 (2022).

71. Ginsburg, K. S. & Bers, D. M. Modulation of excitation-contraction coupling by isoproterenol in cardiomyocytes with controlled SR Ca2+ load and Ca2+ current trigger. Journal of Physiology 556, 463–480 (2004).

72. Ren, S. et al. Implantation of an isoproterenol mini-pump to induce heart failure in mice. Journal of Visualized Experiments 2019, (2019).

73. Burridge, P. W. et al. Chemically defined generation of human cardiomyocytes. Nature Methods 2014 11:8 11, 855–860 (2014).

74. Dhand, A. P. et al. Simultaneous One-Pot Interpenetrating Network Formation to Expand 3D Processing Capabilities. Advanced Materials 34, (2022).

75. Galarraga, J. H., Dhand, A. P., Enzmann, B. P. & Burdick, J. A. Synthesis, Characterization, and Digital Light Processing of a Hydrolytically Degradable Hyaluronic Acid Hydrogel. Biomacromolecules 24, 413–425 (2023).

76. Sala, L. et al. Musclemotion: A versatile open software tool to quantify cardiomyocyte and cardiac muscle contraction in vitro and in vivo. Circ Res 122, e5–e16 (2018).

77. Kim, Y. et al. BeatProfiler: Multimodal in Vitro Analysis of Cardiac Function Enables Machine Learning Classification of Diseases and Drugs. IEEE Open J Eng Med Biol 5, 238–249 (2024).

78. O’Shea, C. et al. ElectroMap: High-throughput open-source software for analysis and mapping of cardiac electrophysiology. Scientific Reports 2019 9:1 9, 1–13 (2019).

79. DI Tommaso, P. et al. Nextflow enables reproducible computational workflows. Nature Biotechnology 2017 35:4 35, 316–319 (2017).

80. Ewels, P. A. et al. The nf-core framework for community-curated bioinformatics pipelines. Nature Biotechnology 2020 38:3 38, 276–278 (2020).

81. Patro, R., Duggal, G., Love, M. I., Irizarry, R. A. & Kingsford, C. Salmon provides fast and bias-aware quantification of transcript expression. Nature Methods 2017 14:4 14, 417–419 (2017).

82. Love, M. I., Huber, W. & Anders, S. Moderated estimation of fold change and dispersion for RNA-seq data with DESeq2. Genome Biol 15, 1–21 (2014).

83. Zhu, A., Ibrahim, J. G. & Love, M. I. Heavy-tailed prior distributions for sequence count data: removing the noise and preserving large differences. Bioinformatics 35, 2084–2092 (2019).

